# Development of Actionable Targets of Multi-kinase Inhibitors (AToMI) screening platform to dissect kinase targets of staurosporines in glioblastoma cells

**DOI:** 10.1101/2022.01.20.477108

**Authors:** Oxana V. Denisova, Joni Merisaari, Amanpreet Kaur, Laxman Yetukuri, Mikael Jumppanen, Сarina von Schantz-Fant, Michael Ohlmeyer, Krister Wennerberg, Tero Aittokallio, Mikko Taipale, Jukka Westermarck

## Abstract

Therapeutic resistance to kinase inhibitors constitutes a major unresolved clinical challenge in cancer and especially in glioblastoma. Multi-kinase inhibitors may be used for simultaneous targeting of multiple target kinase and thereby potentially overcome kinase inhibitor resistance. However, in most cases identification of the target kinases mediating therapeutic effects of multi-kinase inhibitors has been challenging. To tackle this important problem, we developed an Actionable Targets of Multi-kinase Inhibitors (AToMI) strategy and used it for characterization of glioblastoma target kinases of staurosporine derivatives displaying synergy with protein phosphatase 2A (PP2A) reactivation. AToMI consists of interchangeable modules combining drug-kinase interaction assay, siRNA high-throughput screening, bioinformatics analysis and validation screening with more selective target kinase inhibitors. As a result, AToMI analysis revealed AKT and mitochondrial pyruvate dehydrogenase kinase PDK1 and PDK4 as kinase targets of staurosporine derivatives UCN-01, CEP-701, and K252a that synergized with PP2A activation across heterogeneous glioblastoma cells. Based on these proof-of-principle results we propose that application and further development of AToMI for clinically applicable multi-kinase inhibitors could provide significant benefits in overcoming the challenge of lack of knowledge of target specificity of multi-kinase inhibitors.

## INTRODUCTION

Multi-kinase inhibitors (MKIs) and more targeted kinase inhibitors are often used in cancer therapies without exact knowledge of the kinases targeted for the therapeutic benefit (Klaeger *et al*, 2017; Lin *et al*, 2019; Montoya *et al*, 2021; Tang *et al*, 2018). Staurosporines (STSs) are a large family of MKIs originally derived from bacterial alkaloid staurosporine (Gani & Engh, 2010). STSs function as classical ATP mimics and are known to inhibit up to 50 kinases with approximately similar efficiency (Gani & Engh, 2010; Klaeger *et al*., 2017; Tang *et al*., 2018). Regardless of their very wide target spectrum and reputation as “dirty kinase inhibitor” several STS derivatives have reached or has been tested in the clinics. Midostaurin (PKC412) is approved for the treatment of FLT3-mutated acute myeloid leukemia (Montoya *et al*., 2021), whereas another STS derivative UCN-01 (7-hydroxystaurosporine) was tested in phase II clinical trials in metastatic melanoma and relapsed T-Cell Lymphomas (NCT00082017). However, in most cases it is not well established what are the actual kinase inhibitor targets the inhibition of which mediates the therapeutic effects in different indications. Further, in a case of brain tumors STS derivatives are compromised by their pharmacokinetic properties as they do not cross the brain-blood barrier (BBB).

Development of MKIs towards clinical use would benefit from a better understanding of the kinase targets mediating both the therapeutic and potential toxic effects in each disease application. However, generalizable strategies for analysis of actionable MKI targets are currently missing. Here, we present Actionable Targets of Multi-kinase Inhibitors (AToMI) as a generalizable approach to identify actionable co-targets of MKIs. We propose that application of AToMI for clinically applicable MKIs would provide significant benefits in overcoming the challenge of lack of knowledge of target specificity of kinase inhibitors.

## RESULTS

### Strategy for characterization of Actionable Targets of Multi-kinase Inhibitors (AToMI)

Protein phosphatase 2A (PP2A) inhibition drives resistance to several kinase inhibitors in multiple cancer types and thereby PP2A reactivation could be envisioned as a novel therapeutic opportunity to overcome kinase inhibitor resistance (Kauko *et al*, 2020; Kauko *et al*, 2018; Vervoort *et al*, 2021; Westermarck, 2018). Related to glioblastoma (GB), we recently demonstrated strong synergistic activity between PP2A reactivation and clinically tested STS derivative UCN-01 (Kaur *et al*, 2016). However, as UCN-01 targets approximately 50 different kinases at nanomolar concentrations (Gani & Engh, 2010; Klaeger *et al*., 2017) it remains unclear which one(s) of these kinases are involved in synthetic lethality (SL) phenotype observed in combination with PP2A reactivation. To systematically map the UCN-01 co-target interactions relevant to synergy with PP2A reactivation, we devised a functional screening platform consisting of the following steps (Fig. 1):

**Figure 1.**
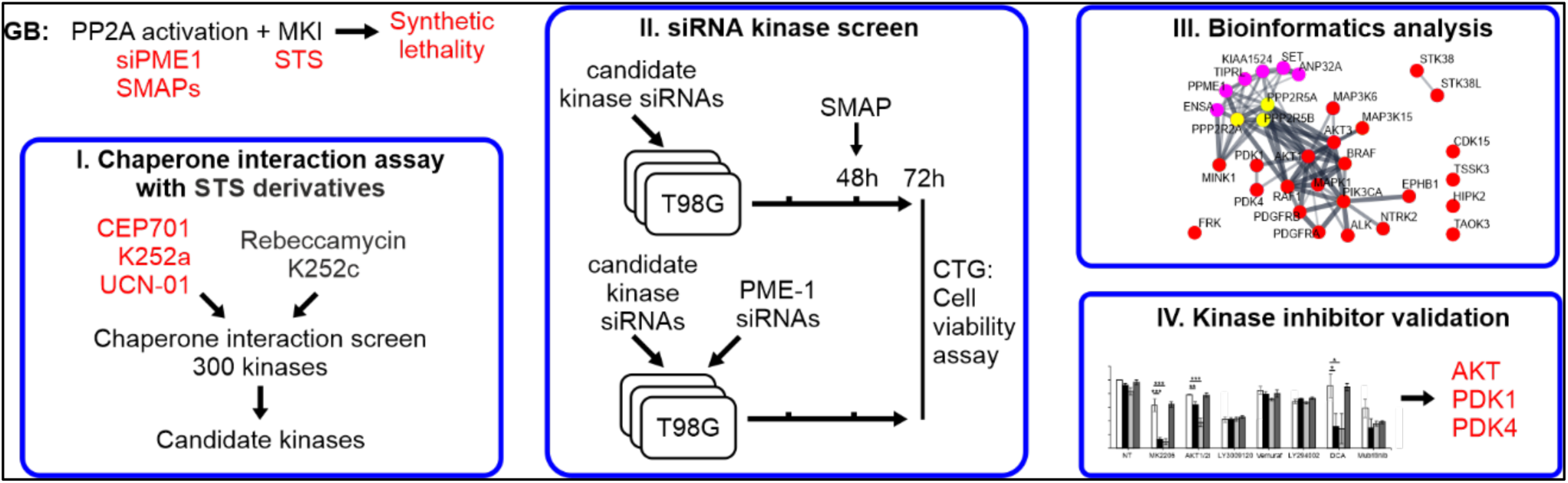
A schematic illustration of AToMI screening platform.

1) Chaperone interaction assay to compare direct kinase binding between UCN-01 and STS derivatives displaying differential synergism with PP2A reactivation in GB cells.

2) siRNA screening for synergistic interaction between PP2A reactivation and targeting of the individual kinase hits from the step 1.

3) Bioinformatics analysis of actionable kinase networks based on steps 1 and 2 for identification of selective small molecule inhibitors for the critical kinase nodes in the network.

4) Small molecule kinase inhibitor validation experiments.

As this strategy could be generally suitable for functional filtering of targets of MKIs, we hereby refer to the screening platform as characterization of Actionable Targets of Multi-kinase Inhibitors (AToMI). The individual technologies used in AToMI are interchangeable with the most suitable technologies for any other application AToMI would be used for.

### Use of AToMI to identify actionable kinase targets of STSs synergizing with PP2A reactivation

By using AToMI, we compared the kinase target profiles of STS derivatives UCN-01, CEP-701 and K252a, previously shown to synergize with PP2A reactivation, against STS derivatives K252c and rebeccamycin that did not synergize with PP2A reactivation (Kaur *et al*., 2016). The differential synergistic activities of these STS derivatives in combination with a small molecule activator of PP2A (SMAPs), NZ-8-061 (Sangodkar *et al*, 2017), were confirmed by colony growth assay in T98G cells (Fig. 2A). In the first step of AToMI, all five STS compounds were screened for their direct kinase protein binding against 300 kinases by Chaperone interaction assay (Taipale, 2018) (Fig. 2B, S1, Table S1). This assay measures the interaction of kinases with their chaperone Cdc37 in the presence (or absence) of kinase inhibitors. Binding of the inhibitor to its target leads to thermodynamic stabilization of the target, which can be detected as weaker interaction between the kinase and Cdc37 (Taipale *et al*, 2013). Using log2 −0.5-fold reduction in chaperone binding as a threshold for interaction, a total of 29 candidate kinases were identified to differentially interact with STS derivatives that synergized with PP2A (CEP-701, K252a, and UCN-01), but not with rebeccamycin or K252c (Table S2).

**Figure 2.**
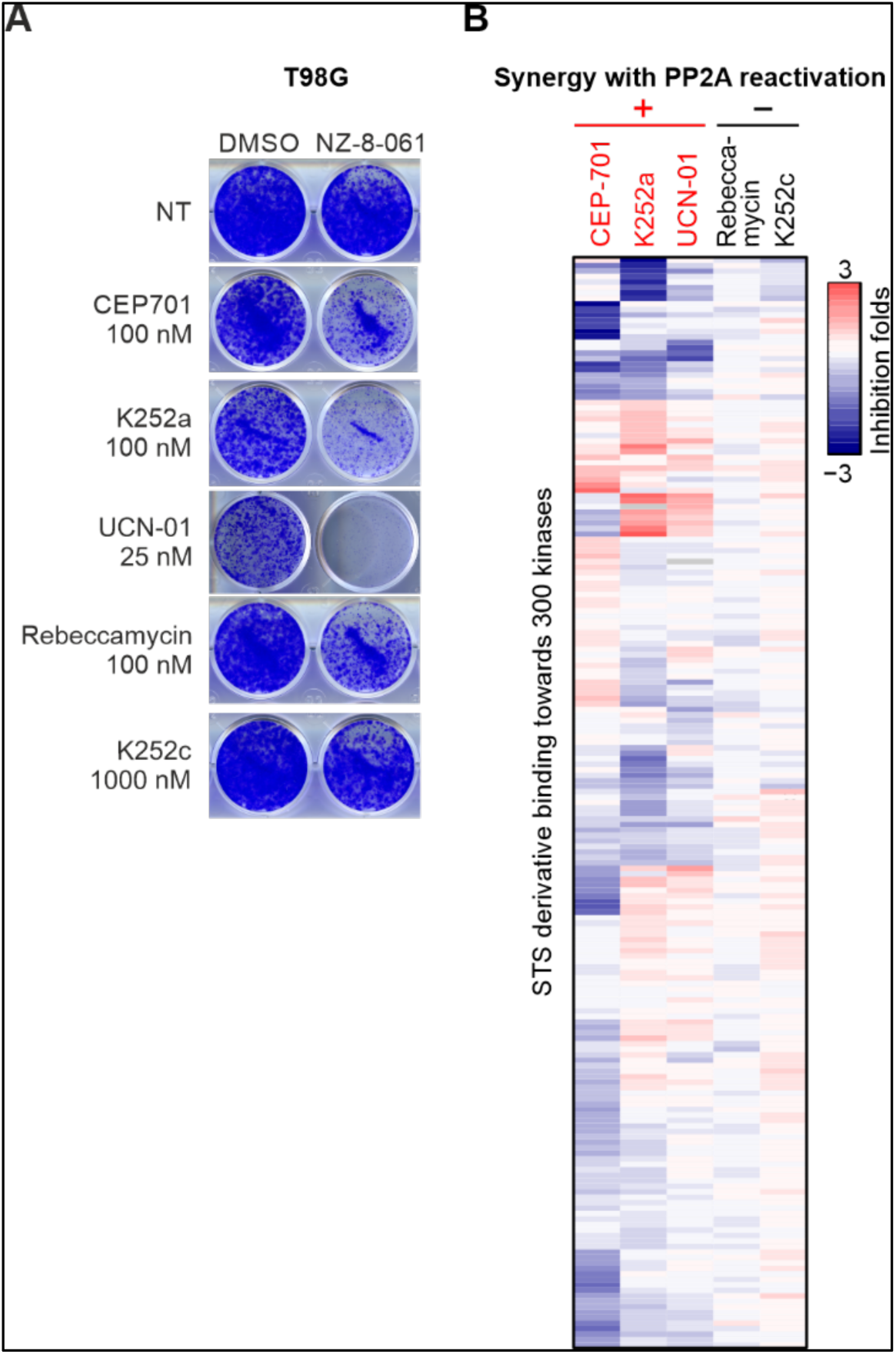
Chaperone interaction assay. **A)** Representative images of colony formation assay in T98G cells treated vehicle (DMSO) and NZ-8-061 in combination with STS derivatives. **B)** Heat map representation of interaction of STS derivatives, CEP-701, K252a, UCN-01, rebeccamycin and K252c, with 300 protein kinases by Chaperone interaction assay. Color scale bar indicates log2 fold changes of kinase/Cdc37 interactions between inhibitor and DMSO treatments.

In the siRNA screening step of AToMI, the goal was to identify among the shared targets of CEP-701, K252a, and UCN-01, individual kinases whose co-inhibition resulted in synergism with PP2A reactivation in cell viability inhibition. The screen was conducted with a custom human kinase siRNA library, which had three non-overlapping siRNAs targeting each kinase. In addition to 29 candidate kinases from the step 1, the siRNA library was extended to include 8 additional kinases frequently altered in GB (Brennan *et al*, 2013; Patel *et al*, 2014) (Table S3). The siRNAs were reverse transfected to T98G cells, and cells were subsequently exposed to PP2A reactivation by NZ-8-061 treatment (Fig. 3A). In the validation screen, we included selected 25 kinases in combination with PME-1 siRNAs to evaluate similarity in drug sensitization between chemical (NZ-8-061) and genetic (PME-1 siRNA) PP2A reactivation (Fig. 3A). The efficacy of PME-1 depletion by tree independent siRNAs was validated by western blotting from parallel samples (Fig. S2). For each kinase siRNA, Gene Activity Ranking Profiles and synergy scores were computed as described in the methods section of siRNA screens. Notably, regardless of the marked differences in the targeting approaches, most of the kinases targeted in both screens were found to synergize both with NZ-8-061 treatment and PME-1 depletion (Fig. 1B), validating both the shared PP2A-induced mode of action, and the broad impact of PP2A activity in kinase inhibitor tolerance in GB.

**Figure 3.**
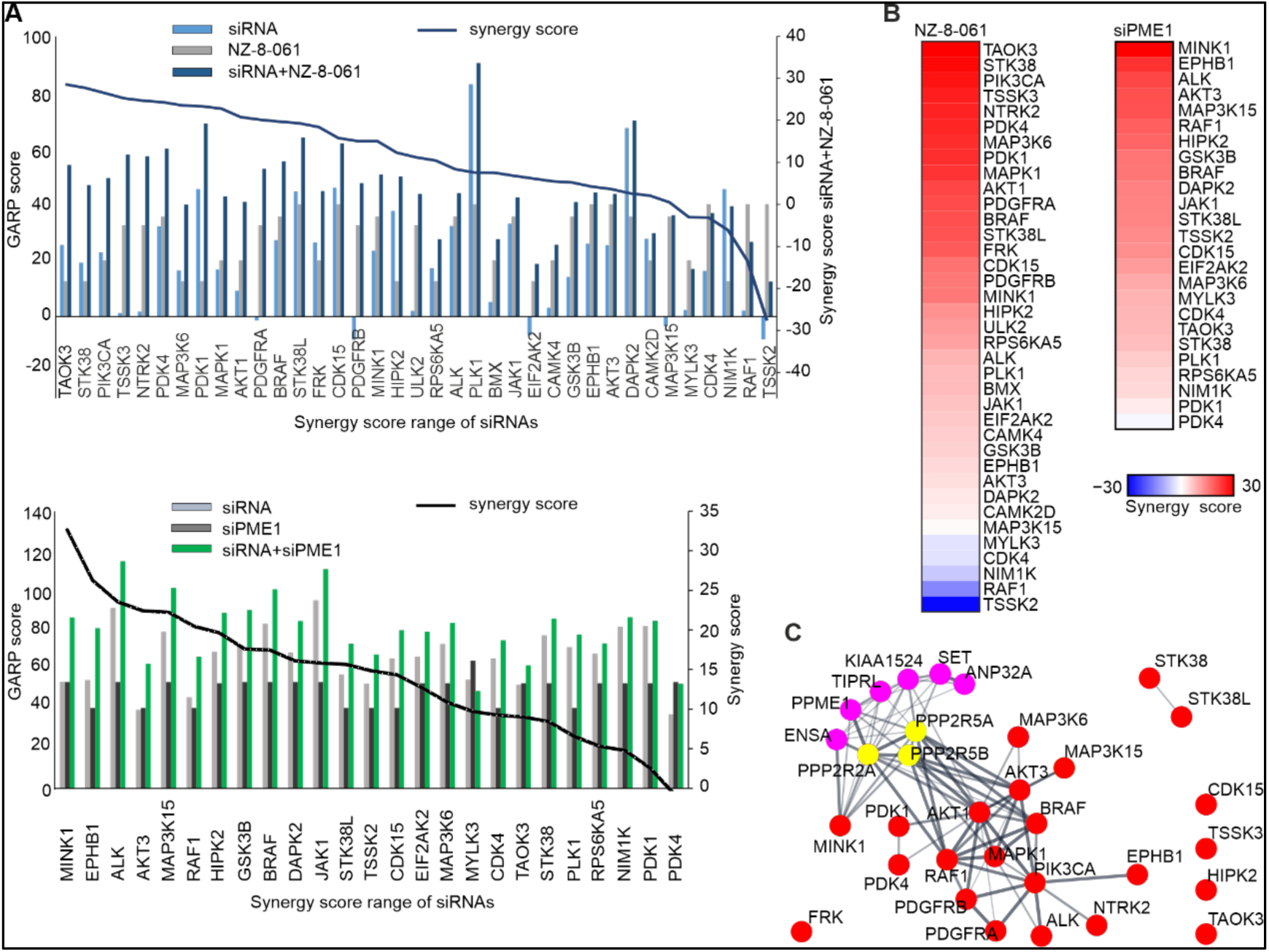
siRNA screening to kinases involved in GB cell synthetic lethality in combination with PP2A reactivation. **A)** GARP scores of siRNA screen in T98G cells under NZ-8-061-treatment or PME-1 depletion (left axis). Kinases were ordered according to synergy scores (right axis). **B)** Heat map representation of kinases involved in synthetic lethality in NZ-8-061-treated and PME-1-depleted T98G cells. Color bar indicates the synergy scores. **C)** STRING interactive mapping of screen kinase hits onto PP2A network.

STRING protein-protein interaction network analysis of the AToMI candidate kinases from the step 2 revealed enrichment of RTK/RAF/MAPK (PDGFR, RAF1, BRAF and MAPK1) and PI3K/AKT/mTOR pathways (PIKCA, AKT1 and AKT3), as well as mitochondrial pyruvate dehydrogenase kinase (PDK1 and PDK4) among the kinases connected to PP2A B-subunits responsible for SL by STS treatment and PME-1 depletion (Fig. 3C) (Kaur *et al*., 2016). As each of these kinase modules were represented also among the kinases that were shared between the NZ-8-061 and siPME-1 synergy targets, we proceeded to testing these GB signaling nodes by selective small-molecule inhibitors. Selectivity of the chosen small-molecule inhibitors was evaluated based recently published target selectivity databases, and for some compounds also by Chaperone interaction assay (Table S4) (Klaeger *et al*., 2017; Tang *et al*., 2018). To facilitate translation of the results, we also considered oral bioavailability and BBB permeability of the compounds in drug selection. The selected 7 kinase inhibitors were screened for cell viability effects in T98G cells with two SMAPs, NZ-8-061 and DBK-1154 (Merisaari *et al*, 2020). As a control, we used an inactive SMAP analog DBK-766, that binds PP2A but is unable to reactivate it even at a concentration of 20 µM *in vitro* (Sangodkar *et al*., 2017). The results show that both NZ-8-061 and DBK-1154 sensitized T98G cells to MK-2206 and AKT1/2i (AKT signaling) (Shariati & Meric-Bernstam, 2019), and DCA (PDK inhibitor) (Michelakis *et al*, 2010; Stacpoole, 2017) used at concentrations that engage their aimed target kinase (Fig. 4A, S3A, B). Importantly, the inactive SMAP (DBK-766) did not synergize with any of these kinase inhibitors (Fig. 4A) and another PDK inhibitor, lipoic acid (Stacpoole, 2017), recapitulated the synergy with SMAPs (Fig. S3C, D). Further validating PP2A reactivation as the mechanism inducing the synergistic drug interaction, also PME-1 inhibition synergized with both MK-2206 and DCA treatments (Fig. S3E). On the other hand, RAF inhibitors (LY3009120 and Vemurafenib), PI3K inhibitor (LY294002), or MINK1 inhibitor (mubritinib) did not display significant combinatorial effect with PP2A reactivation (Fig. 4A).

**Figure 4.**
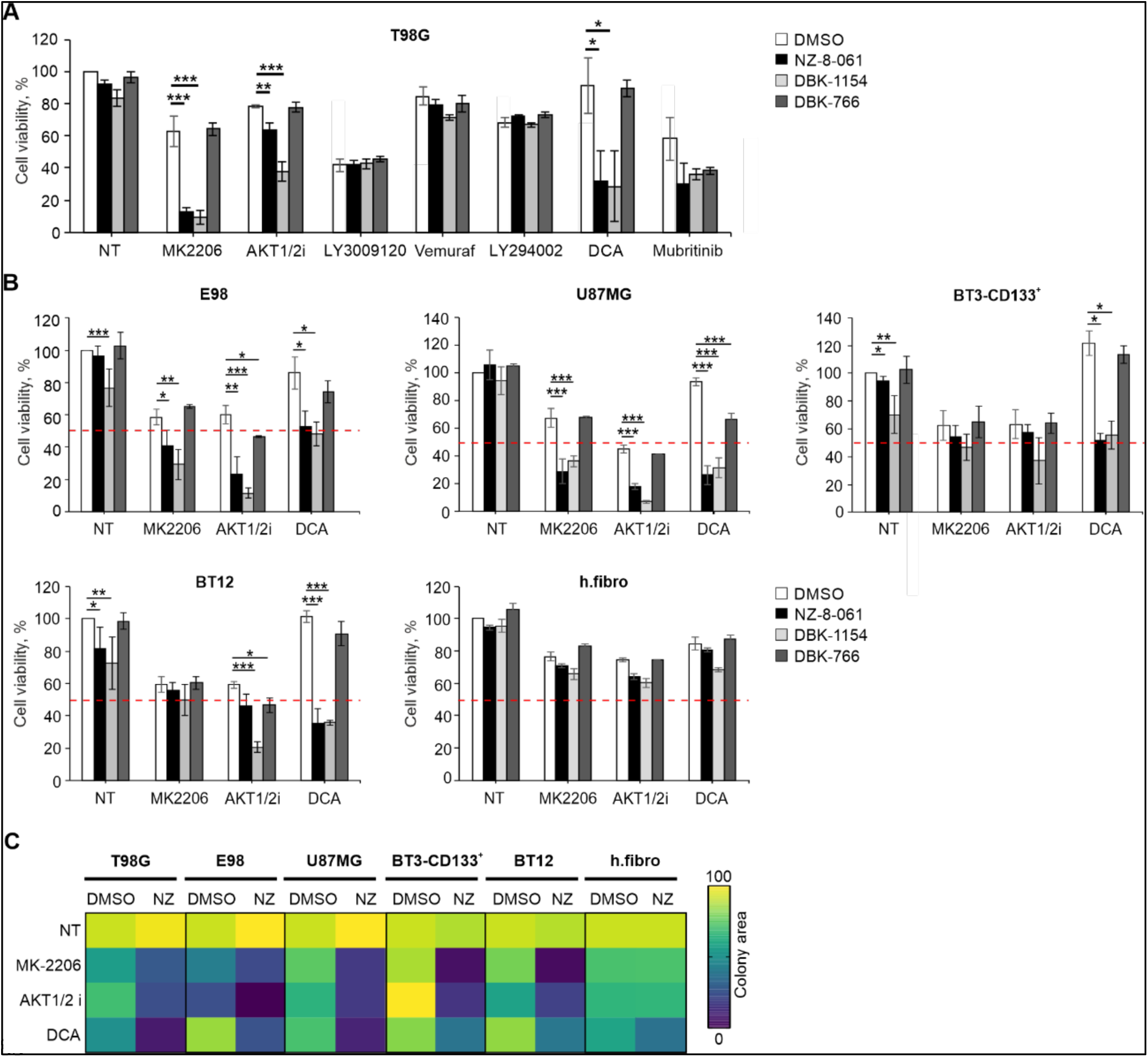
Validation of AToMI results in heterogeneous GB cell lines. **A-B)** Viability of T98G (A) and established GB, E98 and U87MG, and patient-derived GSCs, BT3-CD133^+^ and BT12, cell lines (B) treated with the selected kinase inhibitors alone or in combination with 8 µM NZ-8-061, 6 µM DBK-1154 or 10 µM DBK-766 for 72 h. Human fibroblasts used as a control of normal cells. Data as mean ±SD (n=3 independent experiments). *P<0.05, **P<0.01, ***P<0.001 by Student’s *t*-test. Red striped line indicates 50% inhibition of cell viability which is considered as a cytostatic but not cytotoxic response. **C)** Heat map representation of quantified colony growth assay data in the indicated established GB cell lines, patient-derived GSCs under vehicle (DMSO) or 8 µM NZ-8-061 (NZ) treatment either alone or in combination with indicated kinase inhibitors. Human fibroblasts used as a control of normal cells. (n=2 independent experiments).

Collectively, these results demonstrate the usefulness of AToMI screening for identification of individual actionable target kinases for MKIs.

### Validation of AToMI results in heterogeneous GB cell lines

Cellular heterogeneity and high intrinsic therapy resistance of GS as well as presence of GSCs are major challenges related GB therapies (Gimple *et al*, 2019). We then applied AKT and PDK inhibitors in combination with SMAPs across two additional established GB cell lines, E98 and U87MG, and two patient-derived GSC lines, BT-CD133^+^ and BT12 (Le Joncour *et al*, 2019; Merisaari *et al*., 2020). However, consistently with high intrinsic kinase inhibitor resistance of GB cells (Merisaari *et al*., 2020; Mooney *et al*, 2019), none of the kinase inhibitor monotherapies used at doses that effectively inhibited their intended targets (Fig. S3A, B), induced cytotoxic response (i.e. more than 50% reduction in cell viability) (Fig. 4B). Further, illustrative of the challenge with heterogeneity of GB cell therapy responses, maximal inhibition of cell viability with any doublet combinations was highly variable across the cell lines. In example, BT-CD133^+^ cells were fully resistant to combination of AKT inhibition and PP2A reactivation, whereas the maximal effect of DCA and SMAP combination on E98 cell viability was only 50%. However, indicative of GB cell selectivity of the drug interactions, the human fibroblasts did not show any signs of synergy between kinase inhibition and SMAPs (Fig. 4B). Additionally, the results were confirmed by colony growth assay (Fig. 4C).

## DISCUSSION

MKIs provide an attractive approach to simultaneously inhibit several oncogenic kinases, and some MKIs (e.g., Sunitinib, PKC412), are clinically used as cancer therapies (Montoya *et al*., 2021). However, similar to more selective kinase inhibitors, all tested MKIs have thus far failed in GB clinical trials (Cruz Da Silva *et al*, 2021). STS derivatives targeting more than 50 kinases (Gani & Engh, 2010) could provide a wide enough polypharmacological kinase inhibitor spectrum to target GB driver mechanisms, even in the case of heterogeneous GB cell populations. However, use of STSs as GB therapeutics is compromised by their inability to cross the BBB. To overcome these limitations, and to better understand GB relevant STS target kinases, we developed the AToMI screening platform. As a result of chaperone interaction assay, we found several kinases which selectively bound to STS derivatives. Then by candidate kinases siRNA screens, we identify kinases that synergized with PP2A reactivation by either PME-1 inhibition or by SMAPs. Notably, the kinases which synergized with PP2A reactivation represent the commonly hyper activated pathways in GB. For example, AKT pathway is one of the most dysregulated pathways in GB, and it was well presented in the siRNA screen as depletion of AKT1, AKT3 and PIK3CA all synergized with PP2A reactivation. Another strongly GB associated signaling mechanism was mitochondrial glycolysis, as depletion of both PDK1 and PDK4 synergized with PP2A reactivation. However, AKT and PDK1-4 targeting monotherapies have failed to demonstrate significant survival effects in clinical trials for GB which is consistent with our results that using AKT and PDK-14 inhibitors at doses that inhibit their target kinases has very limited effects on heterogeneous GB cell lines (Dunbar *et al*, 2014; Kaley *et al*, 2019; Stacpoole, 2017; Wen *et al*, 2019). The AToMI approach could clearly identify STS targets that synergize with PP2A reactivation in inhibiting viability of heterogenous GB cells. However, as an additional evidence for high degree of resistance of GB cells towards phosphorylation targeting therapies, even the combinations of PP2A reactivation with either AKT or PDK1-4 inhibitors failed to suppress the viability of most GB cells more than 50% which is yet considered only as a cytostatic effect. Therefore, further studies are needed to explore therapeutic impact of the AToMI identified potential combinatorial approaches in faithful brain cancer models. For this purpose, the AToMI approach was also able to identify more selective and BBB permeable kinase inhibitors with similar biological activity than STS.

Collectively, these results validate the usefulness of AToMI approach for future studies aiming to characterize actionable targets of MKIs in different indications. As the individual technologies used in AToMI are interchangeable with other screening technologies we postulate that AToMI will be widely useful for addressing different biological questions.

## MATERIALS AND METHODS

### Cell culture and reagents

Established human GB cell lines U87MG (gift from Ari Hinkkanen, University of Eastern Finland, Joensuu, Finland), E98-FM-Cherry (gift from William Leenders, Radboud Institute for Molecular Life Sciences, Nijmegen, The Netherlands) and human fibroblasts (gift from Johanna Ivaska, Turku Bioscience, Turku, Finland) were cultured in DMEM (Sigma-Aldrich). T98G cells (VTT Technical Research Centre, Turku, Finland) were cultured in Eagle MEM (Sigma-Aldrich). All growth mediums were supplemented with 10% (except fibroblasts supplemented with 20%) FBS (Biowest), 2 mM L-glutamine and penicillin (50 U/mL)/ streptomycin (50 μg/mL). The patient-derived GSCs BT3-CD133^+^ and BT12 (gift from Pirjo Laakkonen and from Kuopio University Hospital, Kuopio, Finland) were cultured as spheroids in DMEM/F12 (Gibco) and supplemented with 2 mM L-glutamine, 2% B27-supplement (Gibco), penicillin (50 U/mL)/ streptomycin (50 μg/mL), 0.01 μg/mL hFGF-β (Peprotech), 0.02 μg/mL hEGF (Peprotech) and 15 mM HEPES-buffer (Gibco). All cell cultures were maintained in a humified atmosphere of 5% CO_2_ at 37°C.

The following chemicals were purchased from indicated distributors: AKT1/2 inhibitor, CEP-701, sodium salt of dichloroacetate (DCA), lipoic acid, mubritinib, PKC412 and UCN-01 from Sigma-Aldrich; LY3009120 and vemurafenib from SelleckChem; K252a and rebeccamycin from Enzo Life Sciences; K252c from Tocris Bioscience; LY294002 from Calbiochem; MK-2206 from MedChemExpress. Compounds were dissolved in DMSO or mQ (for DCA) and stored at −20°C. SMAPs (NZ-8-061, DBK-794, DBK-1154 and DBK-766) were kindly supplied by Prof. Michael Ohlmeyer (Atux Iskay LLC, Plainsboro, NJ, USA), were dissolved in DMSO and stored at room temperature protected from light.

### Generation of PME-1 knockout T98G cells

*PPME1* deficient T98G cells were generated by CRISPR/Cas9 technology. T98G cells (4 × 10^4^ cells) were plated into 24-well plate and transduced with lentivirus particles containing lentiCas9-Blast plasmid (Addgene #52962). After 18 hours the media were exchanged with fresh media with blasticidin. A single cell-derived clone of T98G/Cas9 was developed, and further transduced with lentivirus particles containing pKLV-PB-U6gPPME1(BbsI)-PGKpuro2ABFP plasmid (gRNA *PPME-1* exon 14: 5’-ACTTTTCGAGTCTACAAGAGTGG, ID 183157785, FuGU, Helsinki, Finland). After 18 hours the media were exchanged with fresh media and 48 hours later the media were complemented with puromycin. Cas9-expression and PME-1 knockout efficiency were evaluated by immunoblot analysis.

### Cell viability assay

Optimized numbers of cells (2.5 × 10^3^ to 5 × 10^3^) were plated onto 96-well plates and allowed to adhere. After 24 hours, cells were treated with vehicle (DMSO) or the indicated compounds. After 72 hours, cell viability was measured using CellTiter-Glo assay (Promega) according to the manufacturer’s instructions using a BioTek Synergy H1 plate reader (BioTek).

### Colony formation assay

Optimized numbers of cells (3 × 10^3^ to 10 × 10^3^) were seeded in 12-well plates and allowed to adhere. The patient-derived GSCs were cultured on Matrigel (Becton Dickinson) coated plates. After 24 h, cells were treated with vehicle (DMSO) or the indicated compounds. After 72 hours, drug-containing media were replaced with non-drug containing medium and incubated until the control wells were confluent. Cells were fixed with ice cold methanol and stained with 0.2% crystal violet solution in 10% ethanol. Plates were scanned and colonies were quantified by ImageJ using Colony area plugin (Guzman *et al*, 2014).

### Chaperone interaction assay

LUMIER (LUminescence-based Mammalian IntERactome) with BACON (bait control) assay was performed as previously described (Taipale *et al*., 2013). In short, 3xFLAG-tagged bait proteins are transfected into 293T cells expressing the Chaperone-Renilla (prey) luciferase in a 96-well plate. After two days, cells are treated with kinase inhibitors (or DMSO) for 1 hour before cell lysis. The cell lysates expressing each bait protein are applied to anti-FLAG coated 384-well plates, which captures the bait protein. The amount of luminescence in the well, after washing off nonspecifically binding proteins, indicates the interaction between the bait protein with the prey protein. After the luminescence measurement, the amount of bait protein is measured with ELISA, using a different, polyclonal anti-FLAG antibody conjugated to horseradish peroxidase.

### siRNA screens

A custom human kinase siRNA library containing three non-overlapping siRNAs targeting each of the 37 kinases was purchased from Qiagen (Table S3). Two independent siRNA screens were done in T98G cells. AllStars negative and AllStars Death (Qiagen) were used as negative and positive controls, respectively. In the first screen, the kinase siRNA library was dispensed in black clear bottom tissue-culture treated 384-well plates (Corning 384) using an Echo acoustic dispenser. The assay plates were used right away or used later in which case they were kept sealed in −20°C until used. For transfection, Opti-MEM medium (Gibco) containing Lipofectamine RNAiMAX (Invitrogen, Thermo Fisher Scientific) was added (5 µL per well) using a Multidrop Combi (Thermo Fisher Scientific) and plates were mixed for 15-30 min at room temperature. After that, T98G (500 cells per well) were added in 20 µL of culture medium using the Multidrop Combi. Final siRNA concentration was 12 nM. After transfection, cells were incubated at 37°C for next 48 hours in the presence of 5% CO_2_. Then cells were treated with NZ-8-061 (5 µM) for the next 24 hours and cell proliferation was measured by CellTiter-Glo (Promega) according to the manufacturer’s instructions using a Pherastar FS plate reader (BMG Labtech). In the second screen, kinase siRNA library in combination with the control (scrambled and AllStars negative) and PME-1 siRNA (three variants, Table S5) was dispensed first as previously described. Then T98G cell were seeded and incubated at 37°C for next 72 hours in the presence of 5% CO_2_. Cell proliferation was measured by CellTiter-Glo. Using collected data for each plate, the following calculations were performed to obtain percentage inhibition values for all wells (% inhibition=100*((averageneg–averagesample)/(averageneg– averagepos))). From the siRNAs targeting the same kinase, Gene Activity Ranking Profile (GARP) score for the kinase was calculated by taking average of two siRNAs with the highest values in inhibition data (or two lowest siRNA values from viability data) (Marcotte *et al*, 2012). Then, the synergy scores for each kinase were computed using Highest Single Agent model (Berenbaum, 1989).

### Western blotting and antibodies

Cell lysates were prepared and separated by SDS-PAGE as previously described (Merisaari *et al*., 2020). Primary antibodies: PME-1 (Santa Cruz, sc-20086, 1:1000), phospho Akt S473 (Cell Signaling, 9271, 1:1000), phospho PDHE1α S300 (Millipore, ABS194, 1:1000), β-actin (Sigma-Aldrich, A1978, 1:10 000) and GAPDH (HyTest, 5G4cc, 1:10 000). Secondary antibodies were purchased from LI-COR Biotechnology and membranes were scanned using an Odyssey Imager (LI-COR Biotechnology).

### Bioinformatics analysis

Cytoscape network analysis software (version 3.9.0) (Shannon *et al*, 2003) was used to visualize the STRING interactive map of hit kinases (Szklarczyk *et al*, 2021). For calculation and visualization of synergy scores, dose-response matrix of NZ-8-061 and UCN-01 combination data were applied to SynergyFinder (version 2.0) web-application (Ianevski *et al*, 2020).

### Statistical analyses

For cell culture experiments, three biological replicates have been performed, and each condition was tested in triplicate, unless otherwise specified. Data are presented as mean ±SD and statistical analyses were carried out using a two-tailed Student’s *t*-test assuming unequal variances. P<0.05 was considered statistically significant.

## Acknowledgements

We thank the High Throughput Biomedicine Unit at the Institute for Molecular Medicine Finland supported by Biocenter Finland. Johanna Ivaska is thanked for useful comments on the manuscript. Taina Kalevo-Mattila is acknowledged for excellent technical support as well as the entire Turku Bioscience personnel for excellent working environment.

## Author contributions

Conception and design: OD, AK, JW. Development of methodology: OD, JM, JW. Experimental work: OD, JM, MT, CS-F, KW. Bioinformatic analysis: LY, MJ. Resources: MO. Writing: OD, JM, TA, JW.

## Data and materials availability

All data associated with this study are present in the paper or the Supplementary Materials.

## Supplementary information

Fig. S1. Interaction of STS derivatives with 300 protein kinases by Chaperone interaction assay.

Fig. S2. Immunoblot assessment of PME-1 in T98G cells siRNA kinase screen samples.

Fig. S3. Hit validation in heterogeneous GB cell lines.

Table S1. Chaperone interaction assay data.

Table S2. 28 kinase hit list.

Table S3. Custom siRNA kinase library.

Table S4. Inhibitor selectivity data.

Table S5. A list of siRNAs.

