## Supplementary figures 1-3 for "Development of Actionable Targets of Multi-kinase Inhibitors (AToMI) screening platform to dissect kinase targets of staurosporines in glioblastoma cells"

### Figure S1

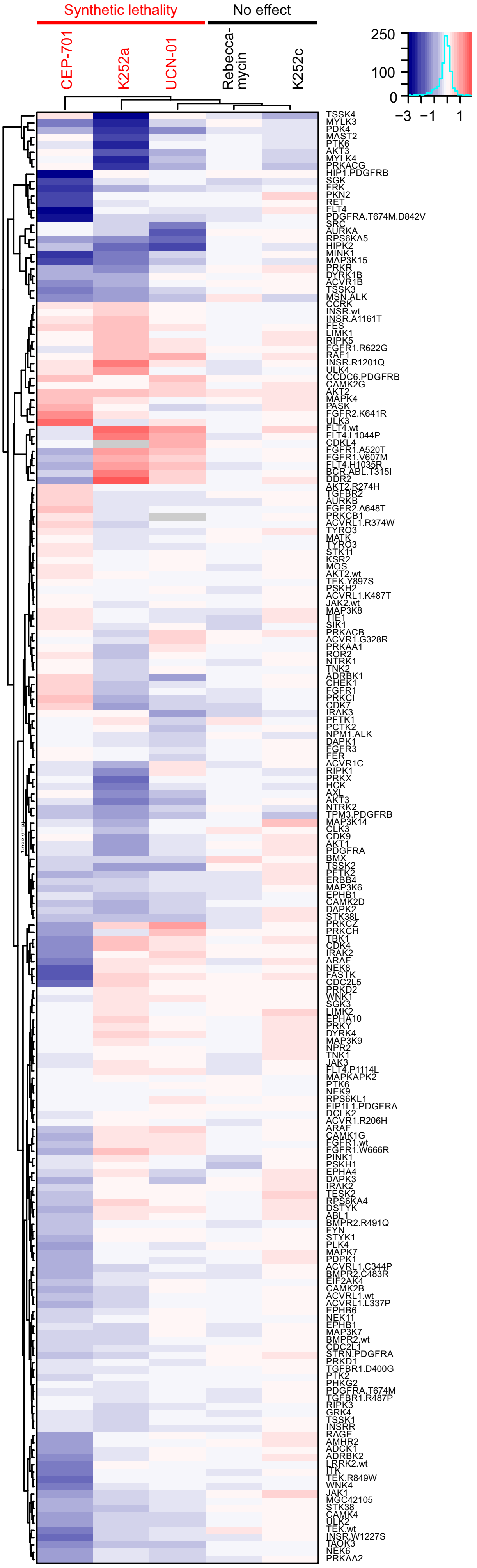


**Figure S1. Interaction of STS derivatives with 300 protein kinases by Chaperone interaction assay.** Heat map representation of interaction of STS derivatives, CEP-701, K252a, UCN-01, rebeccamycin and K252c, with 300 protein kinases by chaperone interaction assay. Color scale bar indicates log2 fold changes of kinase/Cdc37 interactions between inhibitor and DMSO treatments. SL - synthetic lethality inducing drugs (red), NE - no effect inducing drugs (black).

#### Figure S2

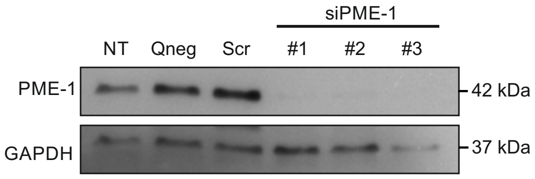


**Fig. S2. Immunoblot assessment of PME-1 in T98G cells siRNA kinase screen samples.**

#### Figure S3

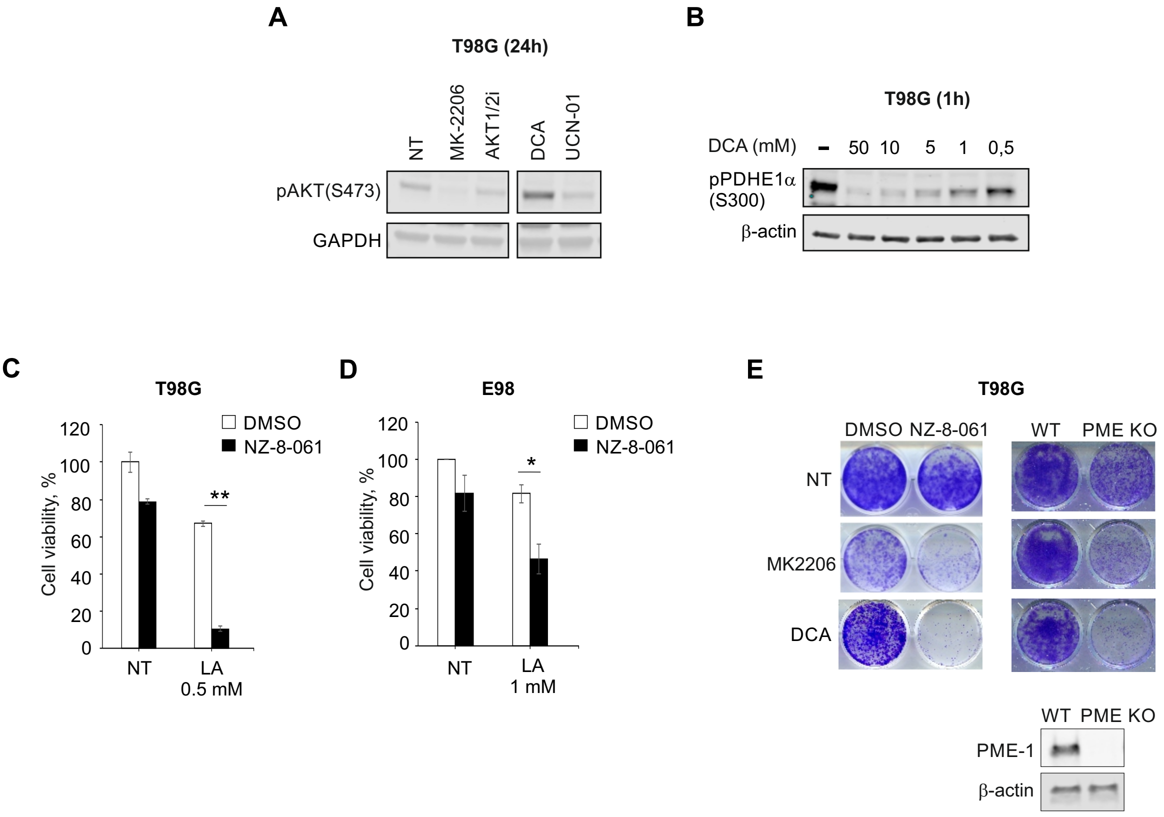


**Fig. S3. Hit validation in heterogeneous GB cell lines. A)** Immunoblot assessment of phosphorylated AKT (S473) after treatment with 5 μM MK-2206, 10 μM AKT1/2i, 20mM DCA and 25 nM UCN-01 for 24 h in T98G. **B)** IImmunoblot assessment of phosphorylated PDHE1α (S300) after 1 h treatment with DCA at the indicated concentrations. **C-D)** Cell viability in T98G (C) and E98 (D) cells treated with NZ-8-061 in combination of lipolic acid (LA) for 72 h. Results are presented as mean ± SD (n = 3 independent experiments). *P<0.05, **P<0.01 by Student's t-test. **E)** Representative images of colony formation assay in T98G cells under PP2A activation by NZ-8-061 (8 μM) or PME-1 knockdown in combination with 5 μM MK-2206 and 20 mM DCA. Western blot analysis of PME-1 knockdown (lower panel).
